# Morphogen And Community Effects Determine Cell Fates In Response To BMP4 Signaling In Human Embryonic Stem Cells

**DOI:** 10.1101/125989

**Authors:** Anastasiia Nemashkalo, Albert Ruzo, Idse Heemskerk, Aryeh Warmflash

## Abstract

Paracrine signals maintain developmental states and create cell-fate patterns in vivo, and influence differentiation outcomes in human embryonic stem cells (hESCs) in vitro. Systematic investigation of morphogen signaling is hampered by the difficulty of disentangling endogenous signaling from experimentally applied ligands. Here, we grow hESCs in micropatterned colonies of 1-8 cells (“μColonies”) to quantitatively investigate paracrine signaling and the response to external stimuli. We examine BMP4-mediated differentiation in μColonies and standard culture conditions and find that in μColonies, above a threshold concentration, BMP4 gives rise to only a single cell fate, contrary to its role as a morphogen in other developmental systems. Under standard culture conditions, BMP4 acts as morphogen, but this effect requires secondary signals and particular cell densities. We further find that a “community effect” enforces a common fate within μColonies both in the state of pluripotency and when cells are differentiated, and that this effect allows more precise response to external signals. Using live cell imaging to correlate signaling histories with cell fates, we demonstrate that interactions between neighbors result in sustained, homogenous signaling necessary for differentiation.

**Summary Statement:** We quantitatively examined signaling and differentiation in hESC colonies of varying size treated with BMP4. We show that secondary signals result in morphogen and community effects that determine cell fates.

## Introduction

Morphogen signaling pathways control cell fate during embryonic development, and can be manipulated to produce particular fate outcomes in human embryonic stem cells (hESCs). During development, all signals both originate from, and are received by, the cells of the embryo, however, cultured cells combine extrinsic influences from the culture medium with endogenous signals passed between cells. In hESCs, secondary signals often perturb the outcome of directed differentiation (Kurek et al., 2015; Warmflash et al., 2014; Yu et al., 2011). Whether endogenous signals are required to maintain particular states, such as the pluripotent state, or to ensure the robustness of differentiation into coherent territories has not been investigated in hESCs. Dissecting the effects of paracrine signals from responses to external stimuli would enable researchers to harness endogenous signals to achieve particular aims, and aid in dissecting the role of these signals in the developing embryo.

The BMP pathway is a conserved morphogen signaling pathway that regulates dorsal-ventral patterning in species from flies to mammals (Bier and De Robertis, 2015) and has also been shown to be essential for mammalian gastrulation (Arnold and Robertson, 2009; Winnier et al., 1995). However, the difficulty in obtaining quantitative data has prevented determining whether BMP functions as a morphogen during mammalian gastrulation. Interestingly, in hESCs, there is increasing evidence that treatment with BMP4 leads to trophectodermal (Horii et al., 2016; Li et al., 2013; Xu et al., 2002) and mesodermal fates (Kurek et al., 2015; Warmflash et al., 2014; Yu et al., 2011), and that the mesodermal fates may be lost when Wnt, Nodal, or FGF signaling is inhibited. Although there is abundant molecular data supporting the identity of these hESC-derived trophoectodermal cells, it has remained controversial (reviewed in (Li and Parast, 2014)) as the correlates of hESCs, the cells of the epiblast, do not give rise to trophoectodermal lineages, and data showing that hESC derived trophoblast cells can function in vivo are also lacking.

When colony geometries are controlled, BMP4 can trigger formation of patterns containing trophectoderm and all three embryonic germ layers (Etoc et al., 2016; Warmflash et al., 2014). These patterns arise in response to homogeneous treatment with BMP4 because of secondary paracrine signals that are required for producing and positioning the mesendodermal territories (Warmflash et al., 2014). Under these culture conditions in which cells are housed within large colonies, it is difficult to disentangle the direct response to the BMP signal from the effects of interactions between the cells (Bernardo et al., 2011). It is therefore unclear whether the different fates induced by BMP4 treatment depend on the dose of BMP4 and, if so, if cells directly read the BMP4 concentration. Quantitative dissection of the cellular response to supplied BMP4 as well as any paracrine interactions that function in the state of pluripotency or during BMP4-mediated differentiation could resolve these important issues.

Here we use a micropatterning approach to isolate the effects of BMP treatment from the secondary endogenous signals that are active both in the state of pluripotency and during BMP-mediated differentiation. To do so, we confined cells to very small colonies ranging from one to eight cells (from here on referred to as µColonies), allowing us to compare isolated cells, which respond only to the exogenous signaling, with cells housed within increasing large colonies where the contribution of paracrine signaling increases. Our results show that, in this context, BMP4 does not act as morphogen but instead functions as a switch and, above a threshold, induces only the trophectodermal fate. In contrast, in standard culture conditions in which colonies may consist of hundreds or thousands of cells, BMP4 elicits both mesodermal and trophectodermal fates in a dose-dependent manner that also requires Nodal signaling and particular cell densities. Further, we find the main effect of secondary signals on the short length scales in μColonies is to enforce a common fate within the colony. This enforcement allows cells to more faithfully remain pluripotent in conditions supporting this state and to differentiate sensitively and homogenously in response to external stimuli. We show that this enforcement is the result of more sustained BMP signaling in larger colonies sizes, and that in standard culture conditions, the outcome of BMP mediated differentiation correlates with the duration of the BMP signal rather than the initial response.

## Results

### BMP4 produces nearly pure populations of trophectodermal cells in μColonies

We first optimized cell seeding such that nearly all μColonies contain between 1 and 8 cells (Fig. 1A,B). Cells in μColonies grown for 42 hours in the pluripotency supporting media MEF-CM expressed the pluripotency markers SOX2, OCT4, and NANOG (Fig. S1 A-C). In the experiments below, we used SOX2 protein expression levels as a marker for hESC pluripotency but show that Nanog obeys similar trends (see Fig. S2). We next assayed the response of μColonies to a range of BMP4 concentrations (0.1-30 ng/ml for 42 hours).

**Figure 1.**
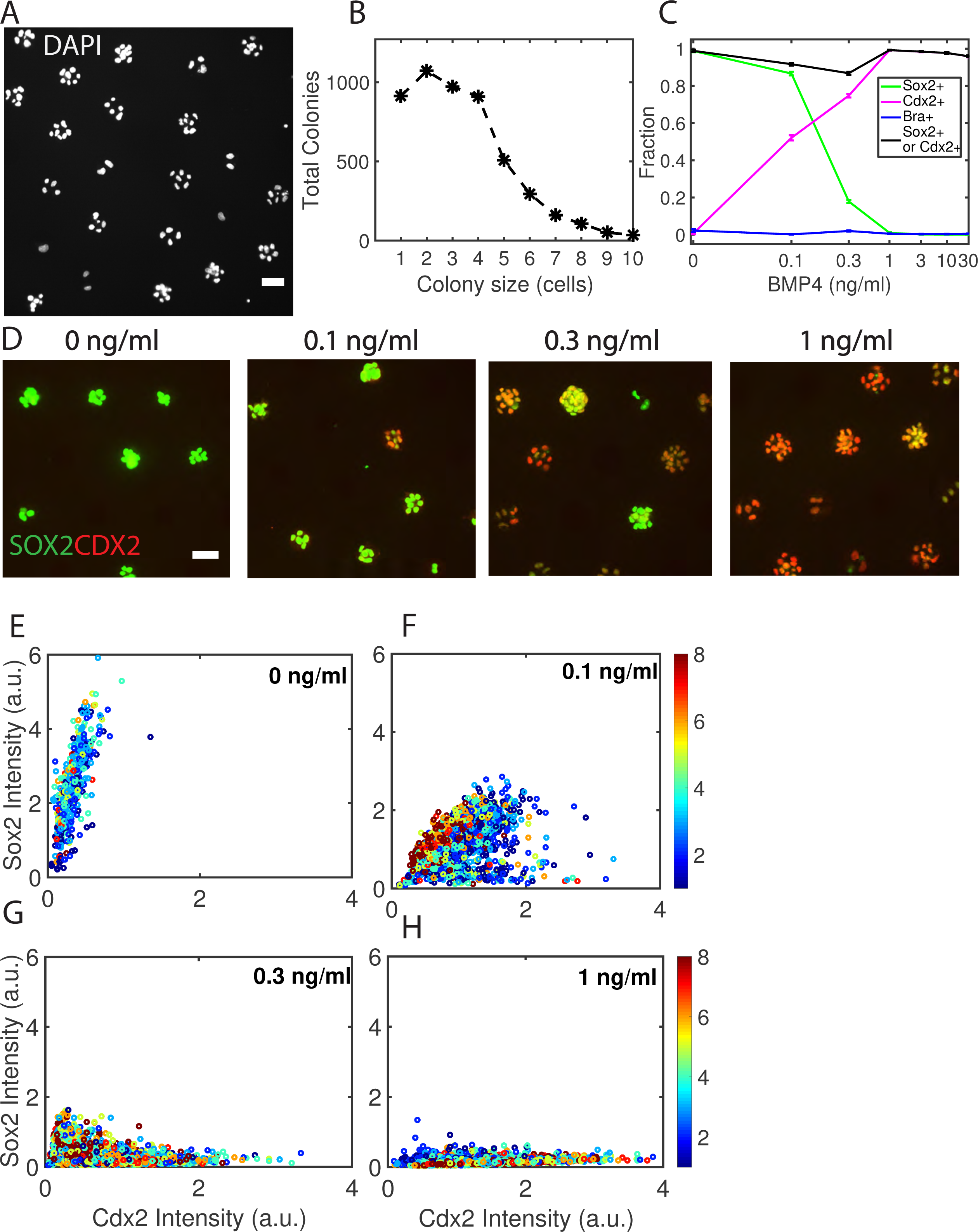
BMP causes differentiation of μColonies to a single fate with a sharp threshold. (A) Representative image of stem cells in μColonies. (B) Representative distribution of colony sizes. (C) Fractions of SOX2, CDX2 or BRA positive cells upon differentiation with BMP4 for 42 hours. (D) Example images of immunofluorescence for CDX2 and SOX2 at the indicated BMP4 doses. (E) Scatter plots of SOX2 versus CDX2 markers for indicated BMP4 concentrations. Each dot corresponds to a single cell while the color code indicates the size of the colony containing that cell. Scale bars 50 μm. See also Fig. S1 and S2.

In response to increasing BMP4 levels, cells within μColonies transitioned from pluripotent (SOX2+) to a differentiated fate expressing CDX2 and GATA3 and lacking expression of BRACHYURY, SOX17, EOMES, NANOG and SOX2 (Fig. S1D-E and Fig. 1C). Consistent with a growing body of literature on BMP4-mediated differentiation (Horii et al., 2016; Li and Parast, 2014; Xu et al., 2002), we identify these cells as trophectoderm, and below we use CDX2 as a marker for this fate. Besides CDX2 and GATA3, all other differentiation markers were detected in less than 2% of cells in the population, and in all conditions, nearly the entire population of cells expressed either the SOX2 marker of pluripotency or the CDX2 differentiation marker. We detected almost no BRA+ cells at any dose (Fig. 1C). BMP4 doses of 0.1 - 0.3 ng/ml produced mixtures of SOX2+ and CDX2+ cells while those at 1 ng/ml or higher yielded nearly pure populations of CDX2+ with complete downregulation of SOX2 expression (Fig. 1D). In contrast, previous literature has shown that larger colonies differentiate to a heterogeneous mixture of fates even in response to much higher doses of BMP4 (Tang et al., 2012; Warmflash et al., 2014). These results establish that cells in μColonies differentiate more sensitively and homogenously than cells in standard-sized colonies in response to BMP4 ligand, and suggest that arrays of small colonies like the ones we employ here may have utility in directed differentiation schemes.

### In standard culture, BMP elicits a morphogen effect that depends on Nodal signaling and cell density

To better understand the lack of mesodermal differentiation in μColonies, we compared the differentiation outcomes in response to a similar range of BMP4 doses for cells grown without confinement to small colonies. We used the pan-mesodermal and primitive-streak marker BRA to determine the extent of mesodermal differentiation. We seeded cells such that the density was homogenous throughout the culture dish and varied this density (see below). We observed a morphogen effect in that the cell fate depended on the concentration of BMP4. Below 2 ng/ml BMP4, cells remained in the SOX2+ pluripotent state, at 2-4 ng/ml cells differentiated to BRA+ mesodermal cells reaching a maximum of approximately 30% BRA positive cells with the remainder CDX2 positive, while at higher doses cells primarily adopted a CDX2+BRA- trophoectodermal fate (Fig. 2A top row, Fig. 2B and Fig. S3A-C).

**Figure 2.**
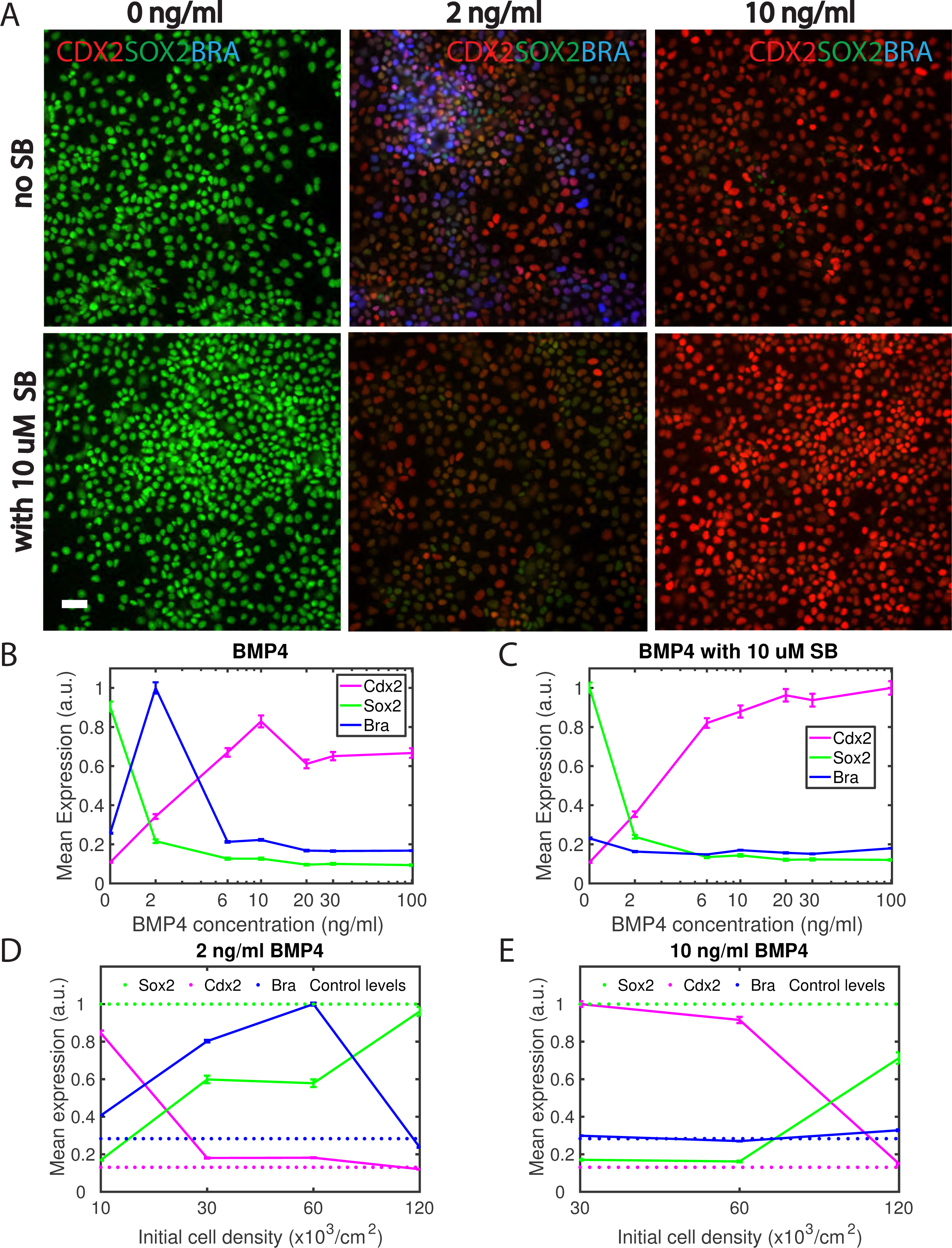
In standard culture, treatment with BMP4 reveals a morphogen effect that depends on Nodal signaling and cell density. (A) Representative images showing SOX2, CDX2 and BRA differentiation in response to two doses of BMP4. Scale bar 50 μm. (B-C) Mean expression of SOX2, BRA and CDX2 markers as a function of BMP4 concentrations with (C) and without (B) 10μm SB431542. The values in (B) and (C) are normalized to the maximum over both sets which were performed in the same experiment. (D-E) Mean expression of SOX2, BRA and CDX2 markers after differentiation with 2 (D) or 10 (E) ng/ml BMP4 with varied initial seeding density. The values in (D) and (E) are normalized to the maximum over the two sets which were performed in the same experiment. Dotted lines represent the levels of expression of the indicated marker under pluripotency conditions. All differentiation experiments were conducted for 42 hours. See also Fig. S3.

If cells directly read the BMP4 concentration, inhibitors of other signaling pathways should not perturb the morphogen effect. We found that treatment with the Activin/Nodal signaling inhibitor SB431542 abolished mesoderm differentiation at all doses so that cells switched between only the SOX2+ and CDX2+ fates as in μColonies (Fig. 2A bottom row, Fig. 2C, Fig. S3D). This supports the idea that the morphogen effect in response to BMP4 requires secondary signals. We next reasoned that the response to secondary signals should be density dependent, and examined the role of cell density in differentiation outcomes. Indeed at the dose of peak BRA induction (2 ng/ml), we only observed BRA-expression at 30 and 60 x 10^3^ cells/cm^2^ but not at lower or higher densities (Fig. 2D, Fig. S3E). At higher BMP4 doses, cells did not express BRA at any cell density but primarily expressed CDX2 at low densities and SOX2 at high densities (Fig. 2E, Fig. S3F). Note that at both 2 and 10 ng/ml BMP4 at high densities, cells failed to differentiate and remained SOX2+, consistent with other reports that BMP signaling and differentiation are inhibited at high cell densities (Etoc et al., 2016). Finally, to explicitly confirm that activating the Activin/Nodal pathway together with BMP stimulation would be sufficient to give rise to mesodermal differentiation, we compared µColonies treated with BMP4 and Activin to those treated with BMP4 alone. Consistent with the above results, we observed substantial mesodermal differentiation in colonies treated with BMP4 and Activin but not in those treated with BMP4 alone (Figure S4). Thus, taken together, our results support a model where only the CDX2 fate is a direct consequence of BMP4 signaling. Mesodermal differentiation can also result at particular doses, but it requires secondary signaling through the Activin/Nodal pathway, and is only induced at particular cell densities. In μColonies treated with BMP-4 alone, cell numbers are likely too low to produce sufficient secondary Nodal signals to induce mesodermal fates, but these can be induced by adding Activin to the media.

### A community effect enforces a common fate within μColonies in both pluripotent and differentiation states

We noted that in the μColony experiments above, even at BMP4 concentrations that produced mixtures of different fates (CDX2+ or SOX2+), the fates of cells within an individual colony were highly correlated, while neighboring colonies often differed in fate, suggesting reinforcement of a common fate within the μColony (Fig. 1D), a phemonenon referred to as the community effect (Bolouri and Davidson, 2010; Gurdon, 1988). To investigate whether a community effect is operating within μColonies, we examined the expression of the SOX2 and CDX2 markers as a function of number of cells in the colony at varying BMP4 doses. Interestingly, under pluripotency supporting conditions, expression of the pluripotency marker SOX2 increased with colony size, while under differentiation conditions, expression of SOX2 decreased with colony size. The differentiation marker CDX2 showed opposite trends: a minor population of spontaneously differentiated CDX2+ cells was observed in 1-cell colonies in pluripotent conditions, and the fraction of CDX2+ cells decreased with colony size. In contrast, the size of the CDX2+ population increased with colony size when differentiated with BMP4 (Fig. 1E-H and Fig. 3A-C). Comparing histograms of expression levels for all cells in the experiment grown in colonies of a particular size under pluripotency conditions, we found a population of cells in 1-cell colonies with reduced SOX2 and enhanced CDX2, and this population was absent in larger colonies (Fig. 3B). This suggests that a fraction of cells spontaneously differentiate to a distinct state and that this differentiation only occurs in colonies with small numbers of cells. We also found similar distributions revealing distinct subpopulations of differentiated and undifferentiated one-cell colonies in differentiation conditions but with the opposite trend: pluripotent cells only persisted in colonies with smaller numbers of cells (Fig. S5A-B). This second population of cells becomes increasingly rare as the colony size increases (Fig. 3C, experimental data). We also confirmed this community effect in a second hESC line (Fig. S5C,D) and that it does not depend on the presence of ROCK-inhibitor in the culture media (Fig. S5E,F).

**Figure 3.**
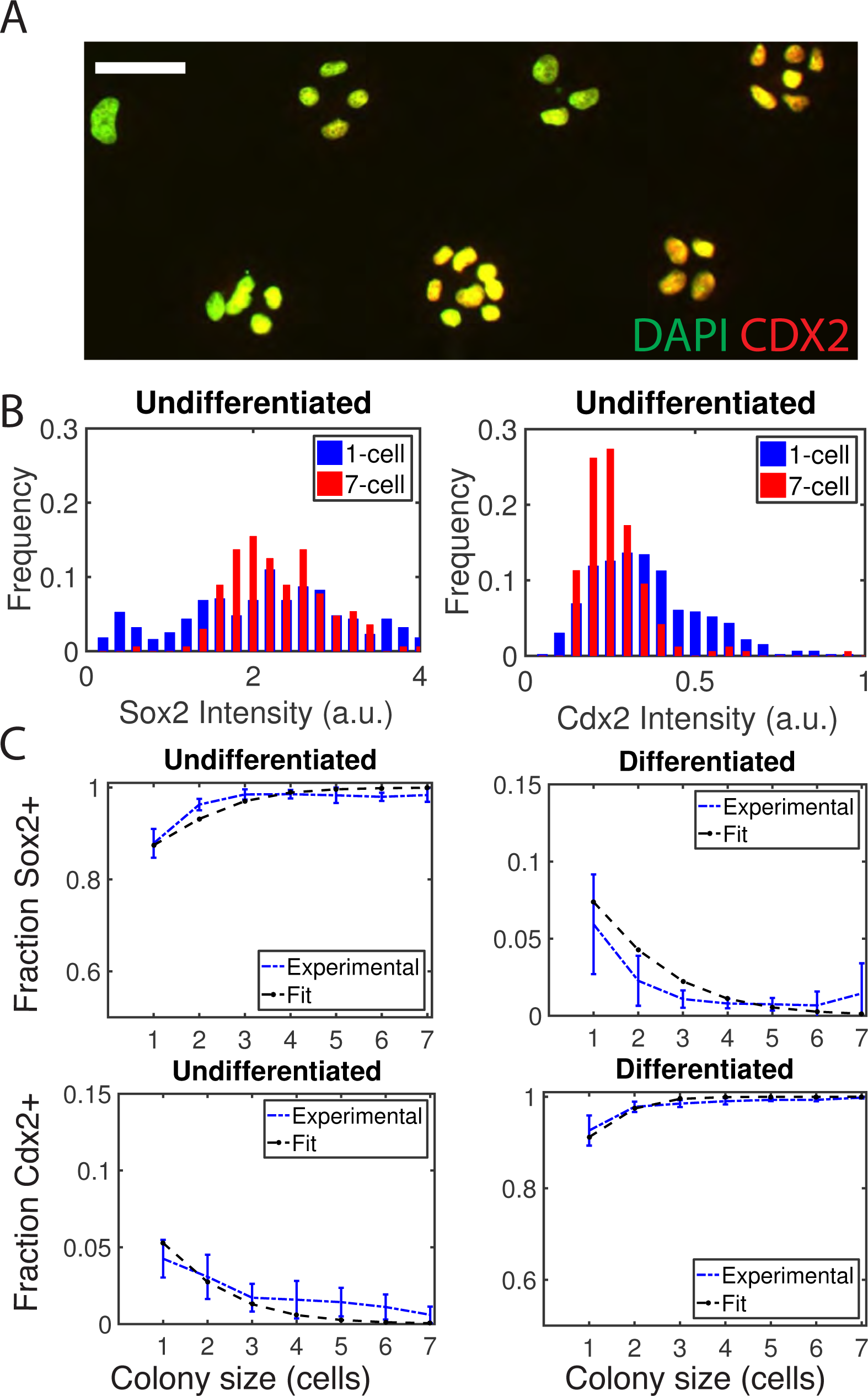
A community effect enforces a common fate in μColonies. (A) Representative image demonstrating the community effect in differentiated conditions (1 ng/ml BMP4). Scale bar 50 μm. (B) Distributions of SOX2 and CDX2 expression in cells of one- and seven-cell colonies in undifferentiated conditions. (C) The fraction of cells expressing a given gene is shown with fits to the Ising-like model. Error bars represent the standard deviation over at least three biological replicates. See also Fig. S5.

### A simple statistical-mechanical model quantitatively accounts for the community effect

The experiments above show that in the μColony system cells can be in one of two states – pluripotent (SOX2+) or trophectodermal (CDX2+). Interactions between cells enforce a common fate inside the colony, while externally supplied BMP4 can bias that common fate towards the CDX2+ state. To explore whether these simple features are sufficient to explain the system’s behavior quantitatively, we exploited an analogy with the Ising model used in statistical physics to describe a two-state system of atomic spins that are coupled to their neighbors and respond to an external field. We made the simplifying assumption that every cell is coupled to every other within a μColony, which is justified by the small colony sizes, and the extensive cell movements we observe in the timelapse experiments below. Within this model, we explored the effects of changing these parameters, and found that increasing either J or the number of cells in the colony will increase the likelihood that all cells in the colony adopt the same fate. As expected, increasing B lead to a general increase in the fraction of CDX2+ cells, with the transition being gradual at low values of J and sharper at high values (Figure S6A-C).

To directly compare the model to data, we performed quantitative fitting of the fraction of CDX2+ and SOX2+ cells as a function of colony size using a separate parameter for the value of B and J at each concentration (see methods and Figure S6). The data for the fraction of cells in each subpopulation as a function of colony size at different BMP4 concentrations was well fit with this simple model (Fig. 3C, black curves). Further, other data not used in fitting the model, such as the distribution of fates within μColonies of a particular size were predicted by the model without further adjustment to the parameters (Fig. S6D). We also examined the fit values of the parameters B and J as a function of the BMP4 concentration (Figure S6E). As expected, the value of B increased with the BMP4 concentration reflecting the increased bias towards CDX2+ fates. Interestingly the value of J was nearly constant reflecting a similar tendency for cells to adopt the same fate within a colony at all BMP4 concentrations. In fact, the data was fit equally well with a model in which the value of J was assumed to be the same for all BMP4 concentrations (Figure S6E). Taken together, these results suggest that within μColonies BMP4 mediated differentiation can be quantitatively explained by only two features – the bias of differentiation towards the trophectodermal fate that increases with BMP4 and the constant coupling between neighboring cells that causes them to adopt the same fate.

### Proliferation rates and clonal composition do not explain the community effect

A simple hypothesis that would partially explain the observed community effect is that some cells are already differentiated upon seeding. If these cells proliferate more slowly, then we would expect colonies that began with differentiated cells would be on average smaller than those containing pluripotent cells. This hypothesis would predict differences in cell cycle as a function of colony size. That is, cells in smaller colonies would be more likely to be arrested in the G1 phase of the cell cycle. To test this hypothesis, we first analyzed the integrated DAPI intensity as a proxy for the total DNA content of the cells, and found that it did not vary with colony size in either pluripotent or differentiation conditions (Fig. 4A). We next created hESCs expressing RFP-Cdt1, a component of the FUCCI system that is expressed only in the G1 phase (Sakaue-Sawano et al., 2008). No differences in the fraction of cells in G1 phase were observed between colonies of different sizes in either pluripotent or differentiation conditions (Fig. 4B), We note that the hypothesis that cell cycle differences underlie the community effect also could not explain our results in the differentiated state where cells expressing pluripotency markers only persist in small colonies.

**Figure 4.**
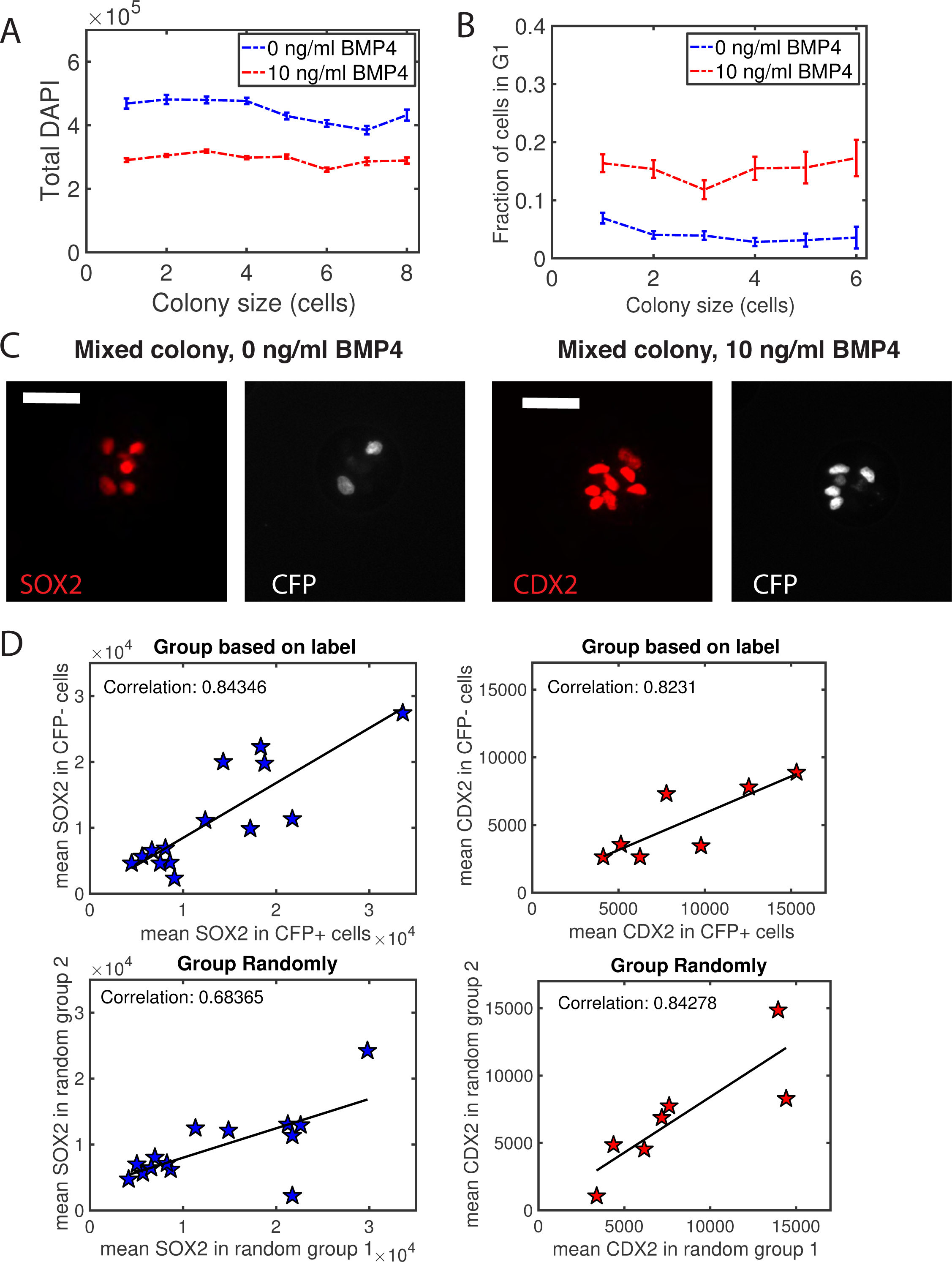
Clonal expansion or cell cycle effects cannot explain the community effect. Mean total DNA content (A) and fraction of cells in the G1 phase of the cell cycle (B) as a function of colony size. In (A) the error bars represent the standard error of the mean calculated separately for each colony size. For (B) the error bars were calculated using bootstrapping method. (C) Representative images of μColonies containing both CFP-positive and CFP-negative cells in pluripotent or BMP4-treated conditions. Scale bar 50 μm. (D) Analysis of mean SOX2 or CDX2 levels in the mixed colonies in pluripotent (blue stars) or BMP-treated conditions (red stars). (Top row) 4-8 cell colonies containing both CFP-positive and CFP-negative cells were identified and the mean SOX2 or CDX2 levels was computed separately for the CFP-positive and the CFP-negative cells. (Bottom row) In the same mixed colonies, the cells were randomly divided into two groups and the mean levels of SOX2 or CDX2 were computed for each group of cells.

To unambiguously establish whether cells within a colony may be more alike because they are clonal derived, we performed an experiment in which we mixed 5% CFP-labeled and 95% unlabeled cells and evaluated their expression of SOX2 in pluripotent conditions or CDX2 in differentiation conditions. If our results can be explained by the clonal composition of colonies, we would expect that most colonies in our experiments are clonally derived, and that in colonies of mixed clones, there are larger differences in expression of markers such as SOX2 or CDX2 between clones than within the cells of the same clone. In larger colonies of 4-8 cells, colonies with CFP-positive cells nearly always had CFP-negative cells as well indicating that multiple clones are typically present despite the uniformity in CDX2 or SOX2 expression (Figure 4C). Moreover, quantitatively, within colonies of mixed CFP positive and negative cells, the mean expression of the CFP positive and negative clones were as correlated as groups of equal numbers of cells in which the cells were chosen randomly without regard to clonal origin (Figure 4D). Thus, these experiments conclusively exclude clonal expansion as an explanation for the uniformity within a colony in either pluripotency or differentiation. Instead, we favor the interpretation that single cells less robustly interpret the supplied signals than small colonies do (see below), and that signaling enforces uniform differentiation within the colony. We also investigated whether the community effect could be affected by modulating the pluripotency-maintaining Activin/Nodal and FGF pathways (Fig. S7A-B) or inhibiting the differentiation-promoting Wnt and BMP pathways (Fig. S7A-B), but we did not observe significant differences in the community effect in any of these cases.

### During differentiation in μColonies, enforcement of sustained signaling underlies the community effect

We next turned to understanding the community effect observed during BMP-mediated differentiation using a reporter cell line for the BMP signaling pathway. We used CRISPR/Cas9 genome engineering to insert GFP at the endogenous locus to form an N-terminal fusion with SMAD4, and isolated a clonal line with a heterozygous insertion of GFP (Fig. S9A-C). Similar fusions have been shown to be faithful reporters of SMAD signaling in the past (Schmierer and Hill, 2005; Sorre et al., 2014; Warmflash et al., 2012). We compared assaying pathway activity with the GFP-SMAD4 reporter and with antibody staining for C-terminally phosphorylated SMAD1/5/8 and found that they give similar dynamics at 1 and 10 ng/ml (Figure S9D-E).

In undifferentiated cells, GFP-SMAD4 localizes to the cytoplasm and translocates to the cell nucleus upon stimulation with BMP4 (Fig. 5A). We performed live confocal imaging beginning approximately four hours before stimulation and quantified the BMP signaling response by measuring the nuclear to cytoplasmic ratio of GFP-SMAD4 during differentiation induced by 10 ng/ml BMP4 (Fig. 5A,B and Movie S1). To increase statistical power, we seeded reduced numbers of cells in μColonies and focused only on the difference between 1 and 2 cell colonies. We note that the designation of number of cells in the colony refers to the number of cells at the time of stimulation. We observe both cell division and cell death in colonies of all sizes, and the final number and the number at the time of stimulation will often differ.

**Figure 5.**
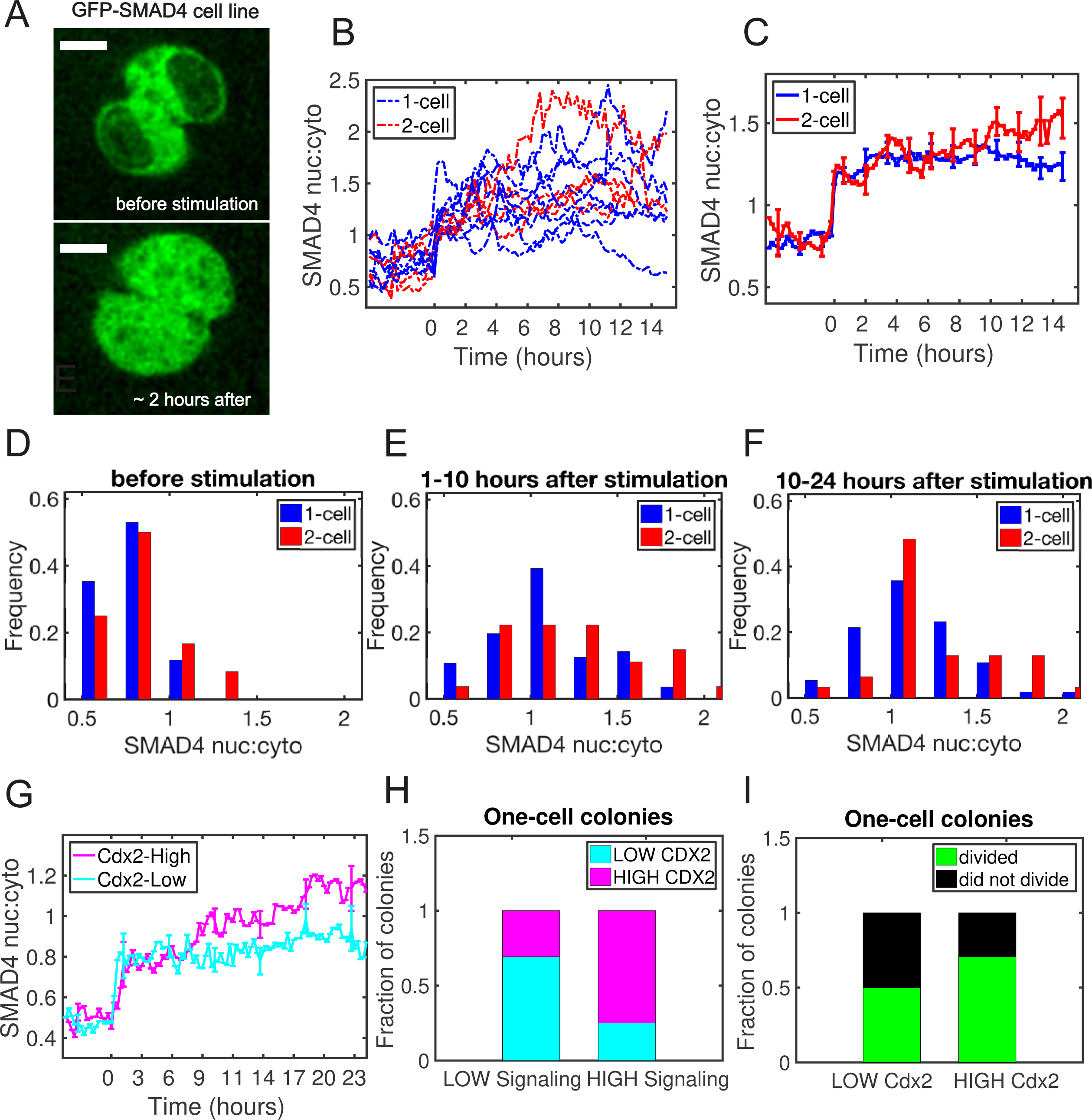
Reinforcement of the BMP signal underlies the community effect in differentiated cells. (A) Representative images of reporter cells. Scale bar 10 μm (see Movie S1). (B) Representative trajectories for the one- and two-cell colonies treated with 10 ng/ml BMP4. (C) Mean signaling trajectories for one and two cell colonies. (D) –(F) Histograms of mean signaling intensity in individual cells over the indicated time intervals. (G) Cells were classified as high or low expressing for CDX2. The mean signaling is shown in each case. (H) Signaling trajectories were similarly binarized as high or low signaling and the fraction of cells with high or low CDX2 examined as function of the signaling level. Due to difficulty tracking individual cells for 42 hours, cells were fixed and analyzed for CDX2 after 24 hours in BMP4. In panels (C) and (H) error bars represent standard error of the mean over trajectories. (I)The same colonies from panel (H) that contained one cell at the time of BMP4 stimulation were grouped depending on whether that cell later divided and then evaluated for CDX2 expression.

The reporter revealed similar signaling intensities in 1 and 2 cell colonies before BMP4 stimulation and in the early response to the ligand up to 10 hours after stimulation. Thereafter, the mean trajectories began to diverge with the two-cell colonies showing higher signaling (Fig. 5C). Examining the distribution of signals in individual cells, we found that this divergence in the mean is mostly due to the presence of one cell colonies that revert to near baseline levels of signaling, while this does not occur in two-cell colonies (Fig. 5D-F). Thus, we hypothesized that cells without sustained signaling will fail to differentiate to CDX2+ cells while the high signaling cells will differentiate.

To test this hypothesis directly, we performed live-cell imaging of one-cell colonies and then fixed these colonies and analyzed their levels of CDX2. We defined cells as low or high signaling depending on whether their temporal average overlapped with the distribution of signaling before stimulation. We found that 75% of high signaling cells but only 31% of low signaling cells differentiated to a CDX2+ cell fate (Fig. 5G). Differences in the mean signaling intensities between CDX2 positive and negative cells became evident after the early phase of response, similar to the differences between one and two cell colonies (Fig. 5H). To see whether the cell cycle might play a role in these results, we also examined whether there were differences in cell division depending on whether cells differentiated to a CDX2+ fate. Dividing and non-dividing cells gave rise to CDX2+ cells in approximately equal proportions (Figure 5I). These data are consistent with a mechanism by which cell-cell interactions serve to maintain the BMP signaling response, perhaps by directly activating the BMP4 gene (Karaulanov et al., 2004; Schuler-Metz et al., 2000), and thereby enforce differentiation to trophectodermal fates. One-cell colonies that lack this reinforcement both signal and differentiate more heterogeneously.

### During differentiation in standard culture conditions, sustained signaling is required for differentiation into CDX2 fate

To investigate the relationship between BMP signaling dynamics and differentiation more generally, we performed dose response experiments under standard culture conditions using the same GFP-SMAD4 cell line. At each dose, we measured the BMP signaling dynamics and then fixed the same cells and analyzed their differentiation to CDX2+ trophectoderm. To avoid the complications of cells adopting multiple fates, we cultured the cells with SB431542 in order to prevent mesodermal differentiation. Interestingly, in the range of 1-10 ng/ml BMP4, the initial response to ligand stimulation was identical and the trajectories only diverged at later time points with 1 ng/ml showing significant decay of the signal and 3 ng/ml showing a small decay as compared to the cells at 10 ng/ml (Fig. 6A,B top panel). These trends were mirrored in the differentiation data, cells at 1 ng/ml largely failed to express CDX2 while those at 3 ng/ml expressed it almost as highly as those at 10 ng/ml (Fig. 6B, bottom panel). Since the initial signaling response was the same in all cases, these data demonstrate that the maintenance of signaling, rather than the magnitude of the initial response, is the determining factor for whether cells will differentiate in response to BMP4.

**Figure 6.**
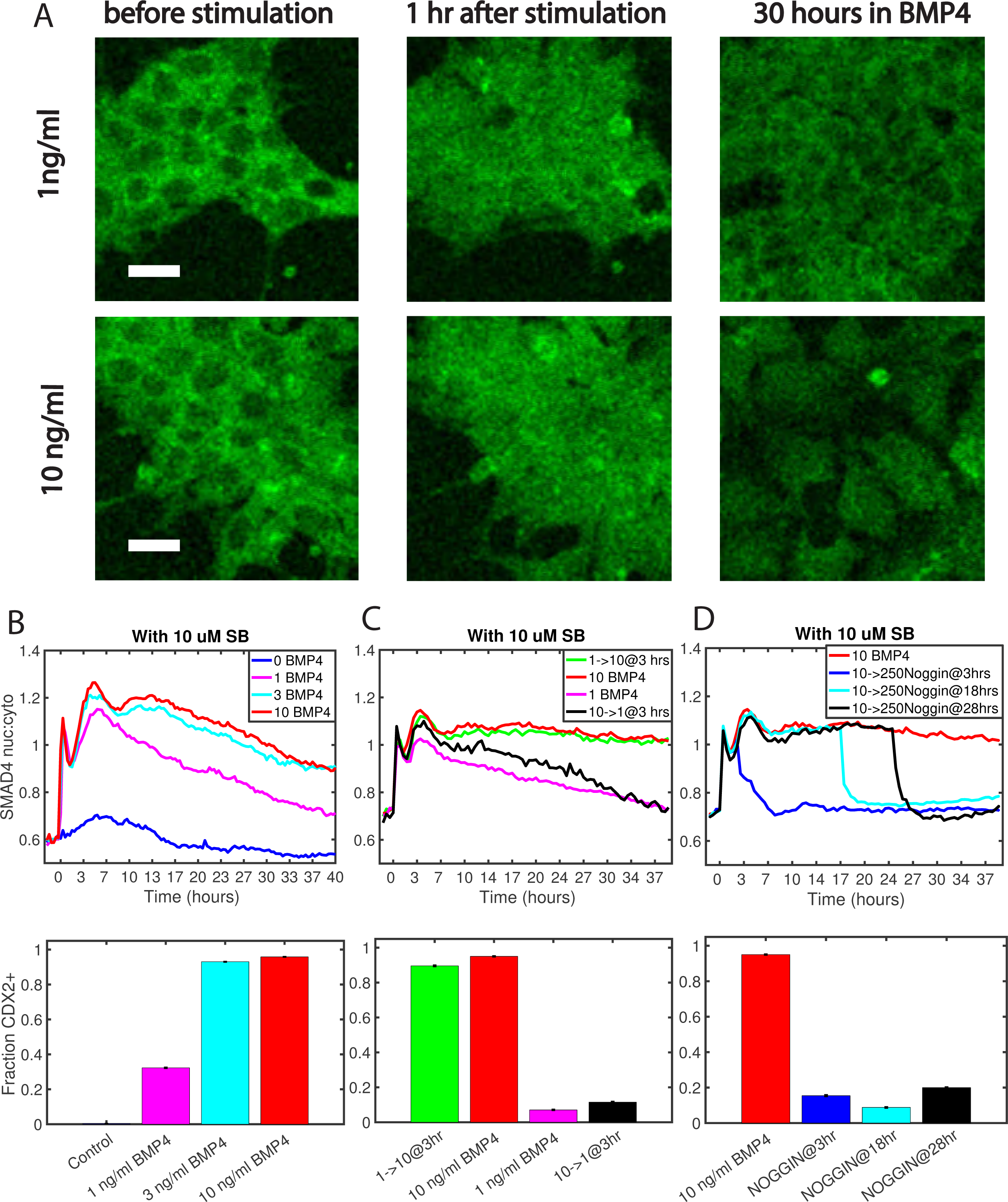
Cells fates correlate with the duration of signaling rather than initial response to BMP4. (A) Representative images of reporter cells in standard culture conditions at the indicated time points following BMP4 treatment. Scale bar 20 μm. Images were acquired every 20 minutes. (B-D) (Top row) The BMP pathway was stimulated with BMP4 or inhibited with Noggin as indicated in the legends. The GFP-SMAD4 reporter line was used to determine the signaling activity as a function of time. (Bottom row) The same cells were then fixed and analyzed for CDX2 expression. In (C), the 1→10 and 10→1 conditions, indicate wells that were treated with 1 ng/ml or 10 ng/ml BMP4 and then the media was swapped between those two wells at 3 hours. In (D), the 10→250 noggin, indicates wells in which the media containing 10ng/ml BMP4 was replaced with media containing 250 ng/ml Noggin at the listed times.

To directly determine whether the initial response or prolonged signaling is responsible for differentiation, we performed experiments in which we treated one group of cells with a high concentration of BMP4 (10 ng/ml) that gave rise to differentiation and another group with a lower one (1 ng/ml) that did not. After 3 hours, we switched the media between these two groups, so that one group was switched from 1 to 10 ng/ml BMP4 while the other was switched from 10 to 1 ng/ml. (Figure 6C). Consistent with a sustained response being required for differentiation, cells initially stimulated with 10 ng/ml and then switched to 1 ng/ml had a decline in signaling and did not differentiate to CDX2+ fates, while those switched from 1 ng/ml to 10 ng/ml had sustained signaling and differentiated. Finally, to directly measure the effect of signal duration on differentiation, we stimulated cells with a high dose of 10 ng/ml BMP4 and then switched to media containing the BMP inhibitor Noggin (250 ng/ml) at variable times after the initial stimulation. As measured by the GFP-SMAD4 reporter, switching from BMP4 containing media to Noggin containing media led to a rapid shutdown of signaling. Importantly, only cells remaining in BMP4 for 42 hours differentiated to CDX2+ fates while those switched to Noggin containing media at 3, 18, or 28 hours failed to differentiate (Figure 6D).

## Discussion

Here we introduce a μColony system that allowed us to separately study exogenous and paracrine signaling in hESCs quantitatively and with cellular resolution. We show that endogenous signals enforce a common fate within the colony both in pluripotent conditions and when differentiated with BMP4. This enforcement of a common fate allows larger μColonies to respond more robustly to signals supplied in the growth media: sustaining pluripotency in pluripotency supporting media and differentiating sensitively and homogenously in response to the extrinsic differentiation signal. We show that under standard culture conditions, BMP4 acts as a morphogen, inducing different fates in a concentration dependent-manner, while in μColonies it switches cells from pluripotent to a single fate, trophectoderm, when supplied above a threshold. This apparent discrepancy is due to the need for secondary signals to produce the morphogen effect in standard culture conditions, and μColonies do not reach sufficient densities to produce these secondary signals. We developed a mathematical model which shows that the detailed statistics regarding the number of cells in the pluripotent or trophectodermal fate as a function of colony size can be predicted from only two parameters: the strength of the bias towards the trophectodermal fates by BMP4 and the strength the interactions between cells that enforce a common fate.

The enforcement of a common fate and greater sensitivity to external signals was observed in the induction of Xenopus animal cap cells to muscle fates by vegetal cells by Gurdon who termed this phenomenon the “community effect” (Gurdon, 1988). This work showed that individual animal cap cells inserted between two pieces of vegetal tissue failed to differentiate, in contrast to larger aggregates that were induced to muscle fates. This suggested that interactions between the animal cap cells are required to robustly interpret the mesoderm differentiation signals emanating from the vegetal cells. In hESCs, cells at higher density have been shown to better maintain pluripotency upon the withdrawal of pluripotency-maintaining cytokines, also supporting the existence of a community effect promoting this state (Peerani et al., 2007). A related observation has been made regarding the levels of Oct4 and Nanog in colonies of mESCs grown in “ground state” conditions (Muñoz-Descalzo et al., 2012): the levels differ between colonies but are highly similar between cells in the same colony suggesting reinforcement of common levels through cellular communication. In theoretical work, Bolouri and Davidson proposed that positive feedback of a signal upon its own transcription could underlie the community effect and applied this idea to the maintenance of the oral ectoderm of the sea urchin embryo through induction of *nodal* gene expression by Nodal signaling (Bolouri and Davidson, 2010). Another theoretical study also found that positive feedback on the signal was sufficient to explain the community effect, and suggested that additional negative feedbacks must operate to prevent the entire tissue from converting to a single fate (Saka et al., 2011). Similarly, in this study, we find that the enforcement of sustained BMP signaling by interactions between the cells is necessary for ensuring that all cells within the colony adopt the same trophectodermal fate.

During development, the community effect serves to ensure a common fate over relatively short length scales, and thereby creates coherent territories of a single cell type. Previous work in hESCs has shown that as colony size is increased, cell-fate patterns emerge (Etoc et al., 2016; van den Brink et al., 2014; Warmflash et al., 2014). It is likely that the community effect plays a role in ensuring the coherence of local territories, but other phenomena must emerge on longer length scales to create these patterns. Future work on embryonic patterning with stem cells can probe this transition to understand the emergence of self-organized patterns.

Cells in µColonies of sufficient size differentiate homogenously in response to very low concentrations of ligand. Here, concentrations of 1 ng/ml induced nearly pure populations of CDX2+GATA3+ trophectoderm, whereas in larger colonies, nearly 100 fold greater concentrations induce a mixture of different fates (Tang et al., 2012; Warmflash et al., 2014). Thus, µColonies seeded at appropriate densities may represent a platform for sensitive and robust directed differentiation.

Our results here suggest that only trophectodermal fates are directly induced from epiblast cells by BMP4, and that it does not directly induce multiple fates in a dose-dependent manner. Experiments with inhibiting secondary signals, modulating cell density, and comparing μColonies to standard culture, establish that there is an apparent morphogen effect in treating hESCs with BMP4, but that this is indirect, relying on secondary signals and only operating at particular cell densities. The role of BMP4 in initiating gastrulation and mesendoderm differentiation both in vivo (Winnier et al., 1995) and in vitro (Bernardo et al., 2011; Kurek et al., 2015; Warmflash et al., 2014; Yu et al., 2011) requires other signals and was not seen in our experiments at any BMP4 dose in µColonies. Our data suggest that that μColonies do not contain sufficient cell numbers to initiate the secondary signals such as Nodal and Wnt that operate during gastrulation in the mammalian embryo (Arnold and Robertson, 2009), and are important for patterning pluripotent cells in vitro (Berge et al., 2008; Warmflash et al., 2014).

It will be interesting to use the methods established here to examine whether these other developmental signaling pathways function directly as morphogens. In vivo evidence from genetic perturbations suggests that Nodal signaling induces multiple different fates in a dose-dependent manner during gastrulation (Dunn et al., 2004; Robertson, 2014), and the µColony system could determine whether this is a direct result of cells reading out the Nodal signal or whether other interactions are required. Similarly, as BMP4 has a documented role as a morphogen in dorsal ventral patterning (Ferguson and Anderson, 1992; Tucker et al., 2008; Wilson et al., 1997), it would be interesting to subject these systems to a similar analysis to determine if cells are directly reading the BMP4 concentration in these cases.

## Author Contributions

A.N., I.H., and A.W. designed experiments. A.N. performed experiments. A.N., I.H., and A.W. performed analysis. A.W. supervised research. A. R. created the GFP-SMAD4 cell line. A.N and A.W. wrote the paper.

## Materials and Methods

### Cell Culture

#### Routine Culture

For regular maintenance, hESCs were grown in mTeSR1 in tissue culture dishes coated with Matrigel overnight at 4°C (dilution 1:200 in DMEMF12). Cells were passaged using dispase every 3 days. Cells were routinely tested for mycoplasma contamination. For imaging experiments under standard culture conditions, cells were seeded onto 8- or 4-well imaging slides (ibidi) at densities of ∼63 x10^3^ cells per cm^2^. For density dependent experiments, the densities were varied as indicated.

#### Micropatterned experiments

We used the micropatterning protocol described in detail in (Deglincerti et al., 2016) with adjusted cell numbers. Briefly, micropatterning experiments were performed using HUESM conditioned by mouse embryonic fibroblasts and supplemented with 20ng/ml bFGF (Life Technologies). We will refer to this media as MEF-CM. The day before seeding onto micropatterns, the media was switched from mTeSR1 to MEF-CM. The next day, a single cell suspension was prepared using accutase, and 5.5x10^4^ cells in 2 ml of MEF-CM with Rock-Inhibitor Y27672 (10 μM; StemCell Technologies) were seeded onto the micropatterned coverslip. Custom-patterned glass coverslips (CYTOO) were placed in a 35 mm dish and coated with 2 ml of 5 μg/ml Laminin-521 (LN521, Biolamina) in PBS (with calcium and magnesium) for two hours at 37°C. After two hours, LN521 was washed out via serial dilutions by adding 6 ml PBS and removing 6 ml (6 dilutions). Then the remaining solution was removed entirely, and cells were placed onto the coverslip and incubated at 37°C. After several hours, the media was changed and the growth factors or small molecules added as indicated in the text.

#### Reagents

The following reagents were used to activate or inhibit signaling pathways: BMP4 (R & D systems, dose as indicated in the text), Activin A (R & D systems, 10 ng/ml), Lefty (R & D Systems, 500 ng/mL), SB431542 (Fisher Scientific, 10 μM), PD0325901 (ESI-BIO, 1 μM) LDN-193189 (ESI-BIO, 200 nM), Y27672 (10μM, ESI-BIO) and IWP2 (EMD Millipore, 4 μM). When increasing FGF levels, we used bFGF (Life Technologies, 100 ng/ml).

#### Cell Lines

All experiments in this work were performed with the hESC cell lines ESI017 (ESIBIO) or RUES2 (A gift of Ali Brivanlou, Rockefeller). GFP-SMAD4 cells were made from the parental RUES2 line by using CRISPR/Cas9 genome engineering to fuse a cassette containing a Puromycin resistance gene (PuroR), a t2a self-cleaving peptide, and GFP onto the N-terminus of SMAD4 so that the locus produces both GFP-SMAD4 and PuroR. Subsequently, cells were nucleofected with an ePiggyBac plasmid containing RFP-H2B driven by the CAG promoter and also containing a Blasticidin (Bsd) resistance gene (ePB-B-CAG-RFP-H2B). Cells were selected with 1 μg/ml Puromycin and 5 μg/ml Bsd. The Cdt1-RFP cell line was created by nucleofecting ESI017 cells with an ePiggyBac construct encoding RFP-Cdt1 driven by the CAG promoter and containing a Bsd resistence gene (ePB-B-CAG-RFP-Cdt1). Cells were selected with 5 μg/ml Bsd. The CFP expressing cells were created by nucleofecting ESI017 cells with an ePiggy construct encoding CFP-H2B driven by the CAG promoter and containing a Neomycin resistance gene (ePB-N-CAG-CFP-hH2B). Cells were selected with 200 μg/ml G418.

#### Immunostaining

Coverslips were rinsed with PBS, fixed for 20 minutes using 4% PFA, rinsed twice with PBS, and blocked for 30 minutes at room temperature. The blocking solution contained 3% donkey serum and 0.1% Triton-X in 1X PBS. After blocking, the cells were incubated with primary antibodies at 4°C overnight (see Table S1). The next day the cells were washed three times with PBST (1X PBS with 0.1% Tween20) and incubated with secondary antibodies (AlexaFluor488 cat#A21206, AlexaFluor555 cat#A31570, cat#A21432 and AlexaFluor647 cat#A31571, dilution 1:500) and DAPI dye for 30 minutes at room temperature. After secondary antibody treatment, samples were washed twice in PBST at room temperature. Coverslips were then mounted in Fluoromount-G (Southern Biotech) and allowed to dry for several hours.

#### Imaging

Entire fixed coverslips were imaged using tiled acquisition with a 20X, NA 0.75 objective on an Olympus IX83 inverted epifluorescence microscope. For live cell imaging RUES2-GFP-SMAD4/RFP-H2B reporter cells were seeded on the micropattern as described above and the patterned coverslip was then moved into a holder (CYTOO) to allow for imaging through the coverslip without any intervening material. Images were acquired on an Olympus/Andor spinning disk confocal microscope with either a 40X NA 1.25 silicon oil objective or a 60X NA 1.35 oil objective. Approximately 4 z-planes were acquired at each position every 12-17 minutes. For live cell imaging in standard culture conditions, reporter cells were seeded onto ibidi slides as described above and imaged on an Olympus FV12 Laser Scanning Confocal microscope with a 20X, NA 0.75 objective at time intervals of 20 minutes. We typically acquired 2-4 hours of data before BMP4 stimulation and 20-24 hours for μColonies and 40 hours for standard culture conditions afterwards.

#### Image analysis

Fixed cell experiments utilized large tiled images that were computationally separated in smaller images of size 2048x2048. As the boundaries of these smaller images do not align exactly with the individual images as originally acquired and stitched together by the acquisition software, images presented may derive from 1-4 individual camera acquisitions. Images of fixed cells acquired at 20X on the epifluorescence microscope were segmented using custom software written in Matlab as described previously (Warmflash et al., 2012; Warmflash et al., 2014). Identified cells were grouped into µColonies based on the distance to their neighbors. Cells within 80 microns were considered to be within a single μColony. We visually inspected colony groupings for accuracy. Mean fluorescent intensities for each cell were quantified and intensities for markers were normalized to the mean intensity of the DAPI stain in each cell. All averages are taken over at least 100 cells. Images from live cell experiments were first processed in ilastik (http://ilastik.org) to create nuclear and cellular masks. Custom MATLAB software was used to postprocess these masks to separate touching cells and to quantify both nuclear and cytoplasmic intensities.

Cell-cell communication model. In the conventional Ising Model, the Hamiltonian of the system of atoms in magnetic field B can be written as a sum of energy due to interactions between the neighboring spins and the energy due to magnetic field:

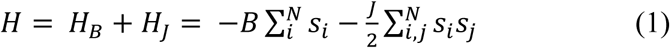

The probability P for the system to be in state σ is given by Boltzman distribution:

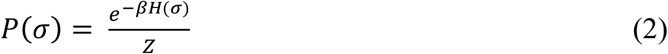

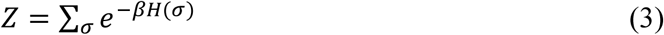

Where Z is the partition function of the system representing the sum of probabilities of all possible states and β is the inverse temperature given by 1/*K_B_*T.

Note from the equations defining the model (1) – (3), that only products βB and βJ appear so that we are free to choose units of energy such that β is equal to 1. We consider a system of size N cells, where the external field B quantifies BMP4 concentration and the parameter J quantifies the strength of the interactions between cells. Since the parameter J > 0, this interaction favors configurations in which neighboring cells have the same identity. We make the simplifying assumption that all cells in the μColony are neighbors which is justified by the small sizes of the μColonies and the extensive cell rearrangements that occur during the observation period (see Movie S1). If we take n cells to be in the CDX2+ state favored by the field B, then (N-n) cells are in the SOX2+ state and the portion of the Hamiltonian due to external ligand is given by:

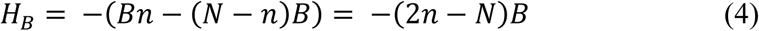

The energy due to cell-cell interactions will be given by:

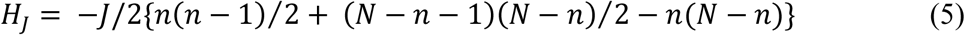

where the first two terms in the sum represent the pairs of interacting cells, that are in the same state (SOX2+ or CDX2+) and therefore contributing to *H_J_* positively. The last term represents the interactions between the SOX2+ and CDX2+ cells and therefore contributes negatively.

The total non-normalized probability for the μColony to have n cells in CDX2+ state and (N-n) cells in SOX2+ state is then:

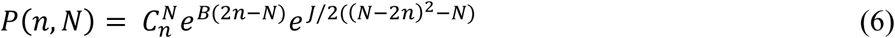

where 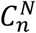 represents the number of combinations of n out of 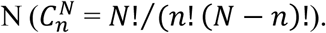 The partition function Z is given by:

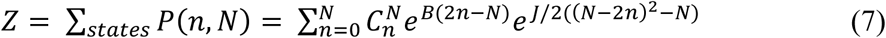

These probabilities are then used to compute averages. For the results in Fig. 3, the average fraction of cells in the CDX2+ state is:

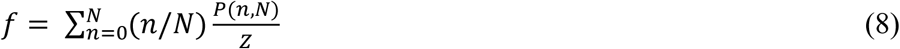

To obtain the theoretical predictions to be compared to the experimental data, we repeat this calculation for all values of N. We then minimized the sum of squares differences between the model predictions and the data using a Monte-Carlo minimization algorithm coded in MATLAB. We performed this fitting independently for each value of the BMP4 concentration, and also performed a fit to all the data in which the value of J was fixed to be the same for all BMP4 concentrations, but the value of B at each concentration was considered a separate parameter (see Figure S6E).

## Acknowledgments

We thank Ali Brivanlou for providing the RUES2-GFP-SMAD4 cell line and the ePB-B-CAG-RFP-Cdt1 plasmid, Eric Siggia and Daniel Wagner for comments on the manuscript, and members of the Warmflash lab for helpful discussions and feedback.

## Data Availability

Custom-written Matlab codes for image segmentation and cell tracking of μColonies can be found on GitHub in repository warmflasha/CellTracker and warmflasha/CellTracker60X.

